# Mind the Outgroup: Influence of Taxon Sampling on Total-Evidence Dating of Pimpliform Parasitoid Wasps (Hymenoptera, Ichneumonidae)

**DOI:** 10.1101/826552

**Authors:** Tamara Spasojevic, Gavin R. Broad, Ilari E. Sääksjärvi, Martin Schwarz, Masato Ito, Stanislav Korenko, Seraina Klopfstein

**Author notes:** Correspondence to be sent to: Tamara Spasojevic, Department of Entomology, National Museum of Natural History, 10th St. & Constitution Ave. NW, Washington, DC 20560, USA.

## Abstract

Taxon sampling is a central aspect of phylogenetic study design, but it has received limited attention in the context of molecular dating and especially in the framework of total-evidence dating, a widely used dating approach that directly integrates molecular and morphological information from extant and fossil taxa. We here assess the impact of different outgroup sampling schemes on age estimates in a total-evidence dating analysis under the uniform tree prior. Our study group are Pimpliformes, a highly diverse, rapidly radiating group of parasitoid wasps of the family Ichneumonidae. We cover 201 extant and 79 fossil taxa, including the oldest fossils of the family from the Early Cretaceous and the first unequivocal representatives of extant subfamilies from the mid Paleogene. Based on newly compiled molecular data from ten nuclear genes and a morphological matrix that includes 222 characters, we show that age estimates become both older and less precise with the inclusion of more distant and more poorly sampled outgroups. In addition, we discover an artefact that might be detrimental for total-evidence dating: “bare-branch attraction”, namely high attachment probabilities of, especially, older fossils to terminal branches for which morphological data are missing. After restricting outgroup sampling and adding morphological data for the previously attracting, bare branches, we recover a Middle and Early Jurassic origin for Pimpliformes and Ichneumonidae, respectively. This first age estimate for the group not only suggests an older origin than previously thought, but also that diversification of the crown group happened before the Cretaceous-Paleogene boundary. Our case study demonstrates that in order to obtain robust age estimates, total-evidence dating studies need to be based on a thorough and balanced sampling of both extant and fossil taxa, with the aim of minimizing evolutionary rate heterogeneity and missing morphological information.

---

The field of molecular dating was established with the first discovery of a correlation between sequence divergence and time (Zuckerkandl and Pauling 1962) and has seen multiple revolutions since then. Today, evolutionary time scales can be estimated using sophisticated computational approaches that incorporate prior knowledge, elaborate mathematical models for sequence evolution and a direct way to integrate time information from fossils.

Until recently, the so-called “node dating” approach (ND) was mainly used for dating phylogenetic trees, with fossils providing minimum ages for specific nodes in a phylogeny. ND requires the prior assessment of fossil placement, which is usually far from straightforward, and it can only incorporate the oldest fossil assignable to a particular node. In addition, one must decide on a probability distribution of the node’s age, because minima are insufficient to date molecular trees; although largely arbitrary, these settings determine the outcome of any ND analysis (Warnock et al. 2011). These issues led to the development of the total-evidence dating approach (TED), which allows inclusion of all available fossils as tips, while accounting for uncertainty in their age and placement on a tree (Pyron 2011; Ronquist et al. 2012a). The downside of TED is that it requires extensive morphological matrices to be compiled, which inform fossil placements and infer associated branch lengths. Some concerns were also raised about the potential lack of clock-likeness of morphological data (O’Reilly et al. 2015), while some authors reported unrealistically old ages from their TED analyses when compared to the oldest known fossils of the group (Beck and Lee 2014; Arcila et al. 2015).

Several methodological modifications to the TED approach have been suggested, such as replacing the initially introduced uniform tree prior (Ronquist et al. 2012a) with the fossilized birth–death tree prior, which models speciation, extinction and fossilization rates to reconstruct branching events, and thus does not necessarily require morphological data (Heath et al. 2014; Ronquist et al. 2016; Zhang et al. 2016). It has also become possible to account for fossils possibly being sampled ancestors (Gavryushkina et al. 2014) and for a diversified sampling strategy of extant taxa (Höhna et al. 2011). Finally, some recent studies combined tip and node dating approaches (O’Reilly and Donoghue 2016; Kealy and Beck 2017; O’Hanlon et al. 2018; Travouillon and Phillips 2018). Unfortunately, these different implementations of TED have often reported disagreeing age estimates (Grimm et al. 2015; Herrera and Davalos 2016; Harrington and Reeder 2017; Kealy and Back 2017; Gustafson et al. 2017), revealing plenty of potential for further improvements of the method (Parins-Fukuchi and Brown 2017; Brown and Smith 2018).

Incorporating more complex models often requires more and better data for improved estimation of model parameters. With the development of next generation sequencing technologies, acquiring large amounts of molecular data is no longer a problem (McCormack et al. 2013; Kjer et al. 2016), but taxon sampling is still a major limiting factor in phylogenetic study design. Numerous simulations and empirical studies have demonstrated positive effects of improved taxon sampling on the estimation of topology (Graybeal 1998; Dunn et al. 2008; Heath et al. 2008; Klopfstein et al. 2017), branch lengths (Fitch and Bruschi 1987; Pick et al. 2010), and parameters of evolutionary models (Zwickl and Hillis 2002; Heath et al. 2008). Nevertheless, thorough taxon sampling is not always easy to achieve, especially in very species-rich groups.

The sampling of outgroup taxa requires special attention: it should ideally include a sufficient sample of taxa that are closely related to, but clearly different from the taxa in question – the ingroup (Wheeler 1990; Nixon and Carpenter 1993; Giribet and Ribera 1998; Graham et al. 2002; Philippe et al. 2011). An inadequately chosen outgroup can significantly affect topology estimates in non-clock analyses by introducing or at least exacerbating long branch attraction (Graham et al. 2002; Holland et al. 2003; Philippe et al. 2011) and/or by increasing compositional heterogeneity of sequences and among-lineage rate variation (Tarrío et al. 2000; Rota-Stabelli and Telford 2008; Borowiec et al. 2017).

Although the influence of outgroup choice on topology estimates is well documented, little is known about its impact on age estimates in molecular dating studies. Some empirical and simulation studies have reported biased age estimates in cases of large among-lineage rate variation (Milne 2009; Wertheim et al. 2012; Duchêne et al. 2014; Soares and Schrago 2015), and highly imbalanced taxon sampling (Linder et al. 2005; Milne 2009; Soares and Schrago 2012, 2015; Duchêne et al. 2015). All of these could be introduced by an outgroup (Heard 1992; Blum et al. 2006; Borowiec et al. 2017). In addition, Duchêne et al. (2015) have shown that the effect of tree imbalance on age estimates is larger when heterochronous sequences are included, as is the case in molecular tip-dating with ancient DNA or in virus studies; a similar effect can be expected when fossil taxa of different ages are included as tips.

In the context of TED, outgroup choice has received little attention, and many studies have employed rather poor outgroup sampling, typically including one to just a handful of outgroup taxa which are severely underrepresented compared to the ingroup taxa, both in terms of morphological characters and fossils (Ronquist et al. 2012a; Arcila et al. 2015; Dornburg et al. 2015; Close et al. 2016; Herrera and Davalos 2016; Kittel et al. 2016; Lee 2016; Bannikov et al. 2017). We here aim to test the influence of outgroup sampling on age estimates in TED, using parasitoid wasps of the family Ichneumonidae as a case study.

The Ichneumonidae is the most species-rich family of parasitoid wasps, with more than 25,000 described species (Yu et al. 2016). We focus especially on Pimpliformes, a monophyletic group comprising nine subfamilies: Acaenitinae, Collyriinae, Cylloceriinae, Diacritinae Diplazontinae, Orthocentrinae, Pimplinae, Poemeniinae and Rhyssinae (Wahl and Gauld 1998; Quicke 2014; Klopfstein et al. 2019). Pimpliformes are especially interesting from a biological perspective, since they cover nearly the entire diversity of hosts and parasitoid strategies known from ichneumonids (Broad et al. 2018). They oviposit into (endoparasitoids) or onto (ectoparasitoids) their host, which they either permanently paralyze (idiobionts) or allow to continue developing (koinobionts), and hosts span almost all holometabolous insect orders, as well as spiders (Araneae). Several attempts have been made in the past to reconstruct the evolution of important biological traits in Pimpliformes (Wahl and Gauld 1998; Gauld et al. 2002; Quicke 2014), but their conclusions were highly dependent on the stability and resolution of the pimpliform phylogeny, which is still in part unresolved (Klopfstein et al. 2019).

Even less well understood than the sequence of events during the radiation of Pimpliformes is their timing and ecological context. The fossil record of ichneumonids is very poorly studied, and most described species come from just a handful of localities (Menier et al. 2004). It starts in the Early Cretaceous with the extinct subfamily Tanychorinae (Kopylov 2010a), but affiliation of this subfamily with Ichneumonidae is somewhat unclear, as its wing venation is intermediate between Ichneumonidae and their sister family Braconidae (Sharkey and Wahl 1992). The Palaeoichneumoninae (Kopylov 2009), also extinct and of a similar age, are thus usually referred to as the oldest ichneumonids (Sharkey and Wahl 1992; Quicke et al. 1999). All remaining ichneumonid fossils from the Cretaceous period have been classified in the extinct subfamilies Labenopimplinae and Novichneumoninae (Kopylov 2010b; Kopylov et al. 2010; Li et al. 2017), except for a single amber fossil that was tentatively placed in the extant subfamily Labeninae (McKellar et al. 2013). The oldest fossil associated with an extant pimpliform subfamily (and an extant genus) is an acaenitine, *Phaenolobus arvenus* Piton, from the latest Paleocene. However, its placement is questionable due to poor preservation, and more reliable records come from two Early Eocene localities, the Green River Formation and Messel Pit (Spasojevic et al. 2018a, 2018b).

No studies have to date attempted to infer the age of Ichneumonidae as a whole or of Pimpliformes in particular. The only previous studies with some bearing on the question are one concerned with the sister family Braconidae (Whitfield 2002) and the other with the entire order Hymenoptera (Peters et al. 2017); both studies included a very sparse sample of ichneumonids. They report considerably different age estimates for the divergence between Ichneumonidae and Braconidae, approx. 138 Ma (Whitfield 2002) and 155–224 (mean 188) Ma (Peters et al. 2017). In addition to obtaining the first age estimates for Ichneumonidae and Pimpliformes and testing the influence of outgroup sampling on these estimates, we also investigate the influence of fossil sampling and placement and discuss the implications of our findings for taxon sampling in dating studies more generally.

## MATERIAL AND METHODS

### Taxon Sampling

We have included 289 tips comprising 210 extant and 79 fossil taxa, with a focus on the pimpliform subfamilies within Ichneumonidae. Among the extant taxa, nine were outside of Ichneumonidae, including seven Braconidae and two more distantly related parasitoid wasps from the superfamilies Chalcidoidea and Evanioidea. The remaining extant taxa included 30 non-pimpliform ichneumonids belonging to 19 subfamilies, and an extensive sampling of Pimpliformes, for which we included 142 of the 188 known genera. For most genera, we included a single representative, but additional species were included in some more heterogeneous genera. The complete list of extant taxa is given in Supplementary file S1.

We aimed to get a good representation of fossil taxa from different time periods: from the oldest ichneumonid fossils from the Early Cretaceous to fossils from the latest Oligocene period (Supplementary file S2). Due to the controversial position of the extinct subfamily Tanychorinae, which is somewhat intermediate in morphology between Ichneumonidae and Braconidae, and poor sampling of the morphological diversity in Braconidae, we excluded the two Tanychorinae fossils from most analyses (but see below).

### Morphological and Molecular Data

We used the morphological matrix from Klopfstein and Spasojevic (2019), but with a strongly expanded taxon sampling. Numerous additional character states were defined to capture the added specimen diversity. The complete morphological matrix consists of 222 morphological characters coded initially for 150 extant and 79 fossil taxa (Supplementary File S3). After observing attraction of fossil taxa to some non-pimpliform ichneumonids for which we had not yet coded any morphological data (“bare-branch attraction”, see Results section), we added another 20 extant taxa to the morphological matrix, increasing the total number of scored extant taxa to 170. Morphology was scored for the same species from which we obtained molecular data, with a few exceptions where we scored a closely related, congeneric species for morphology (Supplementary file S1).

Our molecular dataset includes nine nuclear protein coding genes (primers and protocols in Klopfstein et al. 2019), the D2/D3 portion of the nuclear rRNA gene 28S, and the mitochondrial cytochrome c oxidase (COI). We aimed to amplify these 11 genes for 146 extant taxa and achieved good coverage, with an average of seven genes successfully sequenced per taxon (Supplementary File S4). The sequences were edited and aligned in Geneious 7.1.3 (https://www.geneious.com, Kearse et al. 2012) using the MAFFT v.7.017 plug-in and the algorithm “E-INS-I” (Katoh and Standley 2013). The “translation alignment” option was used for protein coding genes. The complete alignment contained 5423 base pairs (bp) (Supplementary File S5, <TREEBASE link>). The sequences are deposited in GenBank under accession numbers XXX.

### Non-clock Analysis

To assess branch length heterogeneity and to examine the power of our combined molecular and morphological dataset to resolve ichneumonid relationships, we first ran a non-clock analysis. As the analysis showed convergence issues with parameter estimation for most of the 3^rd^ codon partitions of the nine nuclear protein-coding genes and for the entire COI gene, we excluded those from all further analyses. The final alignment thus contained ten genes and 4,011 bp. We partitioned the dataset by gene and codon position and used PartitionFinder 2 (Lanfear et al. 2017) to identify partitions that could be combined (branch lengths = linked, models =all, model_selection = aicc, search = rcluster). In addition, we combined all 1^st^ and all 2^nd^ codon position partitions, respectively, that contained fewer than 30 parsimony informative sites, assuming that substitution model parameter estimates would be very poor for those (Supplementary File S5). All analyses were carried out using Bayesian inference in MrBayes 3.2.6 (Ronquist et al. 2012b) on the HPC cluster UBELIX of the University of Bern, Switzerland (http://www.id.unibe.ch/hpc). The preferred evolutionary model for all partitions was GTR+G+I according to PartitionFinder 2. As MrBayes allows model jumping over the entire GTR subspace, the model parameters for the molecular partitions were set as nst = mixed and rates = invgamma, with all substitution model parameters unlinked across partitions. Morphological characters were analyzed under the Mk model (Lewis 2001), accounting for ascertainment bias (“Mkv”, i.e., only variable characters coded), allowing gamma-distributed rate variation across characters, and ordering all the characters where transition only between neighboring states could be assumed. This model has been identified as the preferred model for the morphological partition in a previous analysis (Klopfstein and Spasojevic 2018). The non-clock analysis included the full set of outgroup taxa (Chalcidoidea, Evanioidea, Braconidae and non-pimpliform Ichneumonidae), with *Gasteruption* (Evanioidea) chosen as the functional outgroup.

We ran four independent runs with four Metropolis-coupled chains each for 150 million generations with a sampling frequency of 1000. The heating coefficient was set to 0.05 in order to increase chain swap probabilities. To summarize the result, we used a conservative burn-in of 50%, while convergence of runs was assessed using typical Markov Chain Monte Carlo (MCMC) diagnostics: the average standard deviation of split frequencies (ASDSF), effective sample size (ESS), and the potential scale reduction factor (PSRF). We also visually inspected the trace plots of the likelihoods and of all parameters for all four runs using Tracer v1.7 (Rambaut et al. 2018).

### Total-Evidence Dating (TED) Analysis

In addition to the settings above, in the TED analysis, we used a uniform tree prior and a relaxed clock model with independent gamma rates (IGR). To set priors on the relaxed-clock model parameters, we relied on the calculations from Ronquist et al. (2012a), putting an exponential prior on the IGR variance with a rate of 37.12, and a lognormal prior on the clock rate with a mean on the log scale of −7.08069 and standard deviation of 1; the standard deviation was decreased compared to Ronquist et al. 2012 to put increased weight on lower clock rates and thus older age estimates, assuming that our dataset contains enough information from the data to correctly estimate the posterior. The prior on the tree age was set to offsetexp(126, 309), with the offset based on the minimum age of the oldest Evanioidea fossil (Deans et al. 2004), while the mean corresponds to the mean age estimate for Hymenoptera from Ronquist et al. (2012a). In the analyses where outgroups were excluded (see below), we used the minimum age of the oldest certain ichneumonid fossils (Palaeoichneumoninae, 112.6 Ma) as an offset. We set a uniform prior on the age of the fossils according to the range of age estimates for the fossil stratum. See Supplementary File S2 for the full list of included fossils and their age intervals with respective references. To obtain the effective prior implied by our settings, including the clock rate (which is only effective when running with data), we performed an analysis without the fossils and thus without temporal information. We ran two independent runs for 100 million generations each and under three different clockrate settings, and always obtained a rather flat age distribution for crown group ichneumonids (Supplementary file S5). Our preferred setting for the clock rate (lognormal with mean of log values = −7.08069 and standard deviation =1), for instance, resulted in a median age estimate of 229.3 Ma and a 95% credibility interval of 51.3 –738.6 Ma (Supplementary file S5). We can thus assume that any more precise age estimates resulting from the analyses with fossils will indeed be informed by the data and not the prior (Parins-Fukuchi and Brown 2017).

It has been shown that relaxed-clock models can lead to topology artefacts, especially close to the root (Ronquist et al. 2012a). We thus set hard constraints on the monophyly of Braconidae, Ichneumonidae and Ichneumonoidea (Braconidae + Ichneumonidae), each of which are widely accepted as being monophyletic and have been recovered in previous analyses (Sharkey and Wahl 1992; Dowton and Austin 1994; Peters et al. 2017), as well as in our non-clock analysis. To improve convergence on the clock rate and tree length parameters, we increased the probability of the respective MCMC moves (MrBayes command blocks are provided as Supplementary File S7). MCMC convergence proved much more difficult to attain than in the non-clock analysis and was thus deemed satisfactory when the ASDSF value was below 0.03, ESS values of all scalar parameters were above 100, and PSRF values were below 1.01. The ESS values between 50 and 100 and PSRF values above 1.01 were still accepted for the tree length, tree height and clock rate parameters in some runs, as it was difficult to get convergence on those even after 150 million generations.

### TED Outgroup Settings

We tested five different outgroup sampling strategies (Table 1): i) “full outgroup” (with all outgroup taxa as in the non-clock analysis), ii) “Braconidae” (all braconids, but excluding the two non-ichneumonoid taxa), iii) “one Braconidae” (a single braconid taxon, *Homolobus*, the only braconid with both molecular and morphological data), iv) “Tanychorinae” (two potentially transitional fossils between braconids and ichneumonids, without including any Braconidae), and v) “Xoridinae” (only members of Ichneumonidae as outgroups). There is strong evidence that xoridines are sister to the remaining ichneumonids (Klopfstein et al. 2019). To set the outgroup, we enforced monophyly of the remaining extant taxa through a topology constraint. None of the fossils were ever included in any topology constraint, but their placement is instead estimated entirely from the morphological data.

**Table 1.**
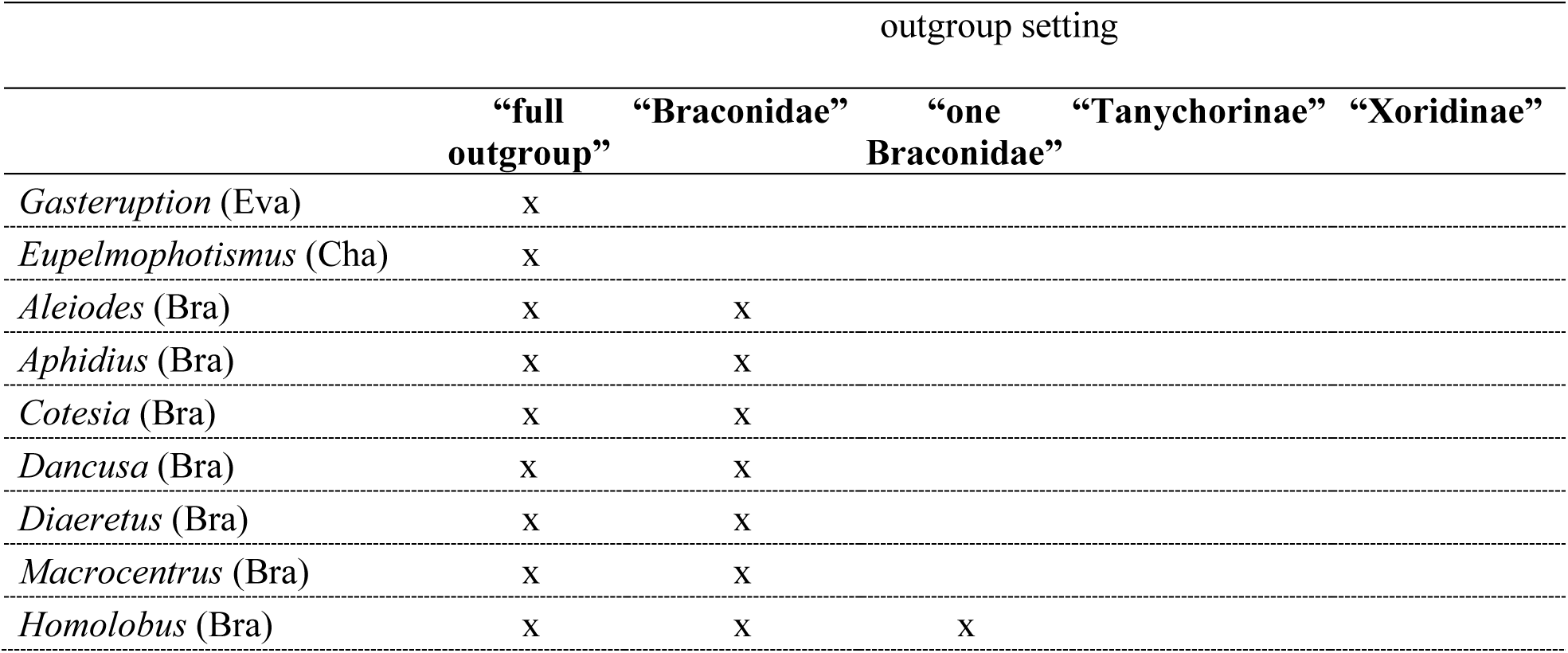
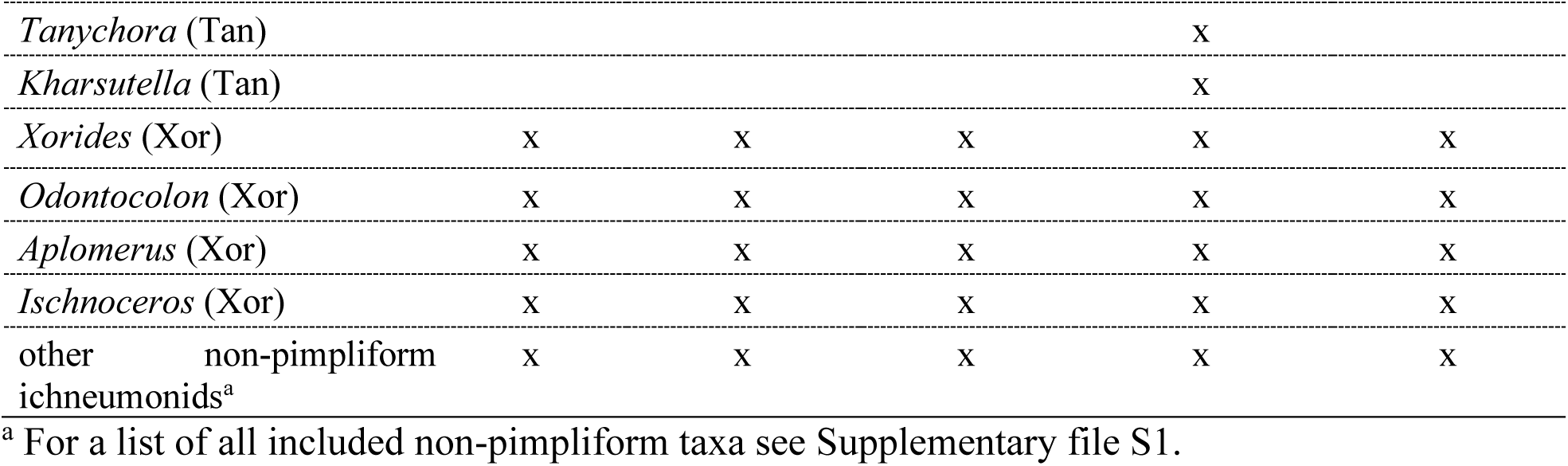
Summary of outgroup sampling strategies. Abbreviations in brackets stand for higher level classification of the outgroup taxa: Eva–Evanioidea, Cha–Chalcidoidea, Bra– Braconidae, Tan–Tanychorinae, Xor–Xoridinae.

|  | outgroup setting |  |  |  |  |
| --- | --- | --- | --- | --- | --- |
|  | “full<br>outgroup” | “Braconidae” | “one<br>Braconidae” | “Tanychorinae” | “Xoridinae” |
| *Gasteruption* (Eva) | x |  |  |  |  |
| *Eupelmophotismus* (Cha) | x |  |  |  |  |
| *Aleiodes* (Bra) | x | x |  |  |  |
| *Aphidius* (Bra) | x | x |  |  |  |
| *Cotesia* (Bra) | x | x |  |  |  |
| *Dancusa* (Bra) | x | x |  |  |  |
| *Diaeretus* (Bra) | x | x |  |  |  |
| *Macrocentrus* (Bra) | x | x |  |  |  |
| *Homolobus* (Bra) | x | x | x |  |  |
| *Tanychora* (Tan) |  |  |  | X |  |
| *Kharsutella* (Tan) |  |  |  | X |  |
| *Xorides* (Xor) | X | X | X | X | X |
| *Odontocolon* (Xor) | X | X | X | X | X |
| *Aplomerus* (Xor) | X | X | X | X | X |
| *Ischnoceros* (Xor) | X | X | X | X | X |
| other non-pimpliform ichneumonids<sup>a</sup> | X | X | X | X | X |

<sup>a</sup> For a list of all included non-pimpliform taxa see Supplementary file S1.

### Fossil Placement

We assumed that an erroneous placement of the oldest included fossils would have the greatest influence, if any, on the age estimates. We thus used “RoguePlots” as described in Klopfstein and Spasojevic (2018) to examine the placement of the Cretaceous fossils on 1,000 evenly sampled trees from the four runs in relevant analyses (R package available at https://github.com/seraklop/RoguePlots). In the “Xoridinae” outgroup setting, the Cretaceous impression fossils were predominantly placed on some terminal branches leading to non-pimpliform taxa without morphological data (see Results section). We thus ran an additional analysis with the “Xoridinae” outgroup sampling scheme and an improved morphological matrix, which now did not contain any outgroup taxa without morphological data.

## RESULTS

### Tree Resolution and Topology

The backbone of the majority-rule consensus trees from both the non-clock and TED analyses was unresolved when fossils were included, which also reflected in the node support values for lower-level relationships. However, when the fossils were excluded from the sampled trees before summarizing them, the resolution at the backbone was strongly improved and all the basal pimpliform nodes were highly supported in the non-clock analysis (posterior probability >0.95). There were only a few topological differences between the non-clock and TED consensus trees, all concerning weakly supported nodes in both of the analyses (Supplementary files S6 and S7).

Most of the expected higher-level relationships outside of Pimpliformes were recovered, such as Xoridinae as the sister group to all other ichneumonids, and the monophyly of the higher groupings Ophioniformes, Ichneumoniformes and Pimpliformes (Fig. 1). Within Pimpliformes, Diplazontinae were recovered as the sister group of the remaining subfamilies. The majority of pimpliform subfamilies were recovered as monophyletic, the exceptions being Cylloceriinae, Diacritinae and Pimplinae (if we disregard a few taxa with alternative placements, see below). The tribe Pimplini was recovered separately from the other pimpline tribes and close to the base of Pimpliformes. It comprised two clearly separate clades, *Xanthopimpla* + *Lissopimpla* and the remaining Pimplini genera, including *Echthromorpha*.

**Figure 1.**
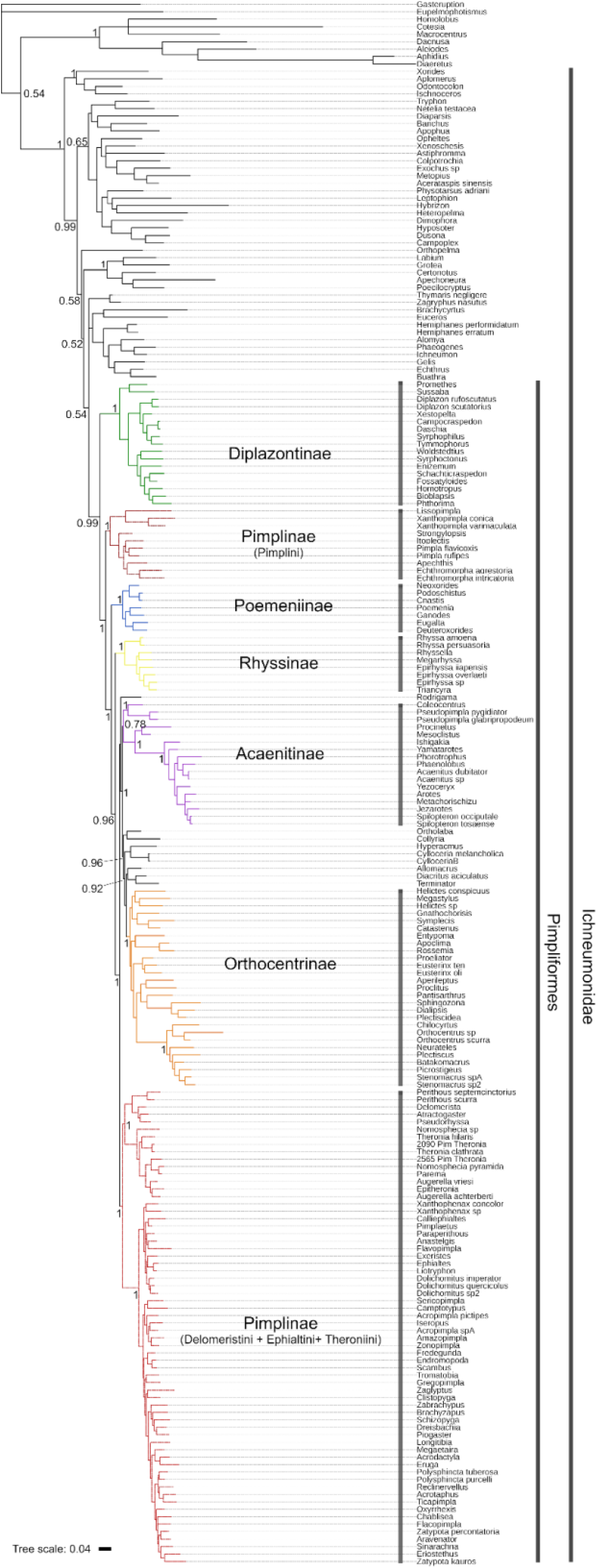
Majority rule consensus tree of the non-clock analysis of the “full outgroup” setting. The tree contains only extant tips, as fossils were excluded prior to summarizing the tree samples from the Bayesian analysis. Posterior probability values are given only for the nodes of interest.

The Poemeniinae genus *Rodrigama* was recovered as the sister taxon to a clade that includes all koinobiont endoparasitoids in Pimpliformes (except Diplazontinae). Within that clade, Acaenitinae had the most basal position, while the relationships among the remaining subfamilies were poorly resolved. The monophyly of Acaenitinae was supported, but *Pseudopimpla* (Ephialtini) clustered with *Coleocentrus* (Acaentinae). For three additional genera, unexpected positions were highly supported: *Ortholaba* (Diacritinae) was recovered with *Collyria* (Collyriinae)*, Rossemia* (Cylloceriinae) grouped with *Apoclima* (Orthocentrinae), and *Terminator* (Orthocentrinae) grouped with *Diacritus* (Diacritinae).

### Impact of Outgroup Sampling on Age Estimates

Both the age estimates and corresponding 95% credibility intervals (CI) were strongly influenced by the outgroup-sampling strategy (Fig. 2). Age estimates were consistently older when outgroup taxa other than ichneumonids or Tanychorinae were included (i.e., more distant and more poorly sampled outgroups) (Fig. 2, Supplementary file S7 and S8). Credibility intervals were widest when Tanychorinae, which only had very few coded morphological characters, were used as outgroup, followed by the strategy with seven braconid taxa. (Fig. 2, Supplementary file S8).

**Figure 2.**
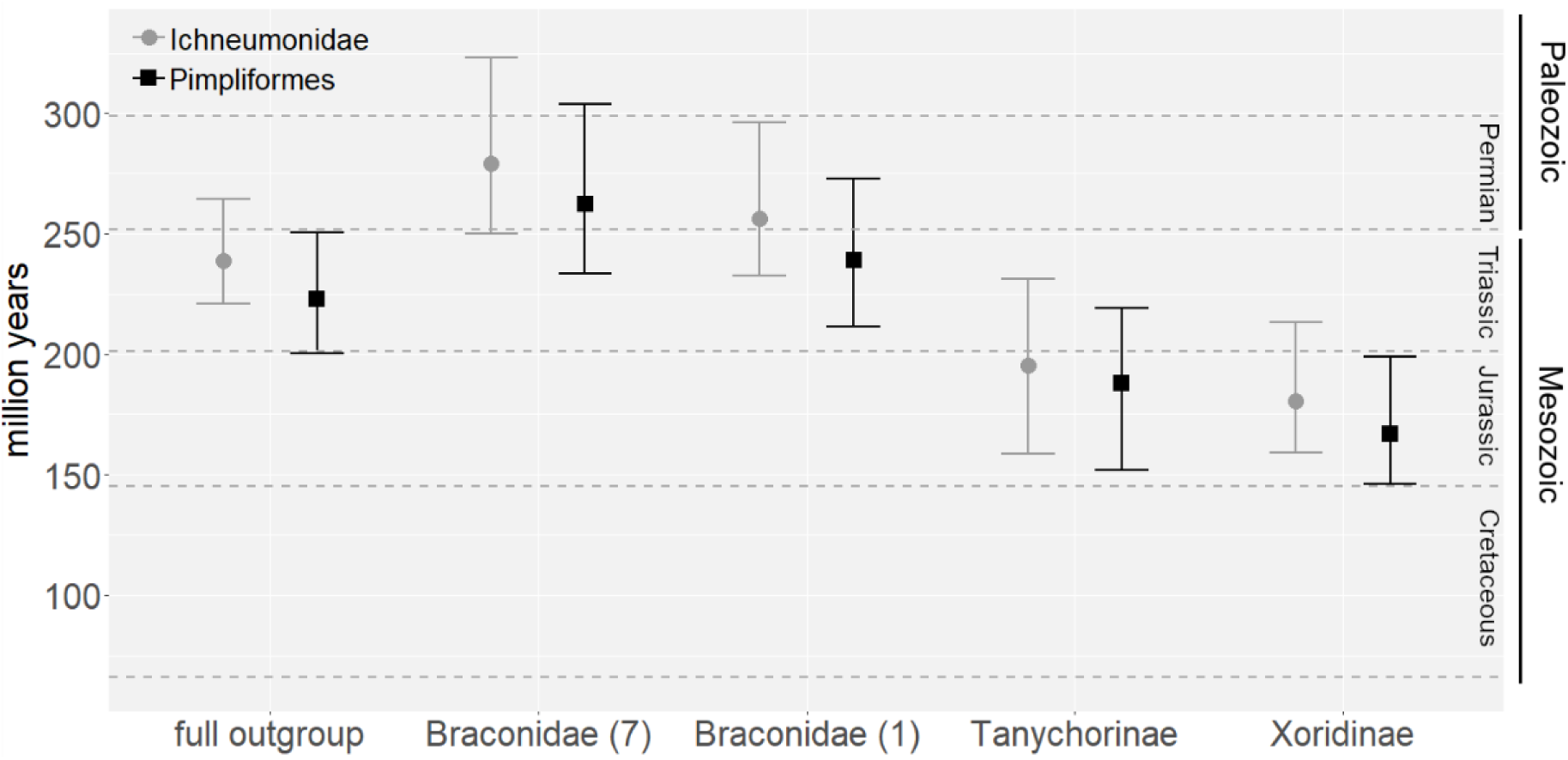
Age estimates in million years for Ichneumonidae and Pimpliformes across different analyses. The median and 95% credibility intervals for different outgroup settings are plotted on the y axis. Numbers in brackets indicate included number of Braconidae taxa. Dashed horizontal lines indicate transitions between major geological periods.

In addition, we assessed the impact of outgroup settings on the variance of the clock rate to detect possible inadequacy of the relaxed clock model to correctly estimate evolutionary rates when distant outgroups are included. The estimated variance for the relaxed clock (IGR) varied considerably across the different outgroup settings (Fig. 3). It was nearly twice as high in the analyses with distant outgroups (“full outgroup” and “Braconidae”) compared to the analyses with close outgroups (“Tanychorinae” and “Xoridinae”). The exception was when a single braconid was included, in which case the estimated clock variance was similar to the analyses with only close outgroups.

**Figure 3.**
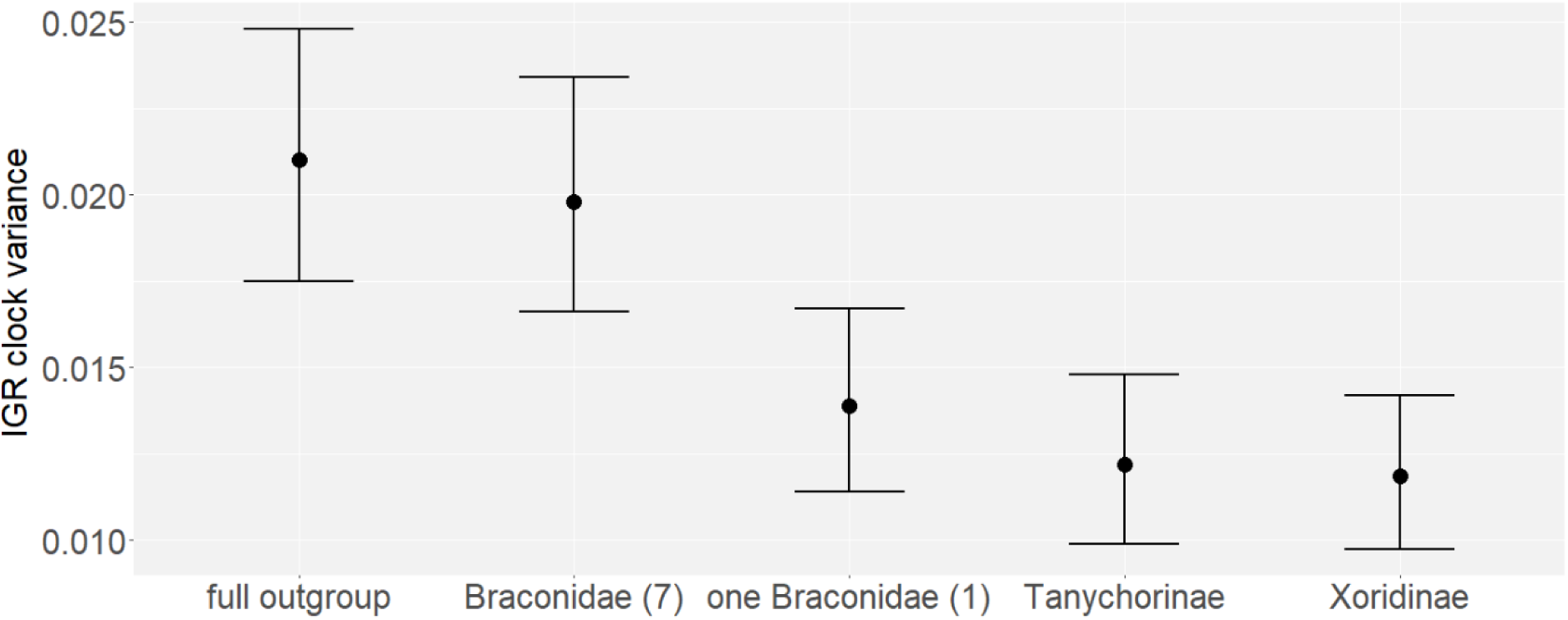
Estimates of the variance of the relaxed-clock. Median and 95% credibility intervals are plotted on the y axis across different outgroup settings. Numbers in brackets indicate included numbers of Braconidae taxa.

### Placement of Cretaceous Fossils

Placements of most of the Cretaceous compression fossils in our initial analyses with different outgroup settings were rather similar: they clustered predominantly within crown-groups of non-pimpliform ichneumonid subfamilies (Fig. 4a, Supplementary file S8). The highest placement probabilities were centered around extant Banchinae (*Banchus* and *Apophua*) and Tersilochinae (in the case of Labenopimplinae and *Tryphopimpla xoridoptera*) and/or on the branch leading to *Orthopelma* (especially in Palaeoichneumoninae), with only a few weakly supported placements in other parts of the tree (less than 10% attachment probability). All Labenopimplinae were placed with the highest probability (34–44%, respectively) on the branch leading to *Banchus*. The overall support that Labenopimplinae belong to crown group Banchinae was quite high (46–53%), and it was even higher if we also consider stem Banchinae (63–73%). *Tryphopimpla xoridoptera* was mostly associated with *Apophua* (34%) with a total probability of attachment within Banchinae of 55% and within extant or stem Banchinae of 71%. In contrast, the small undescribed ichneumonid (3311_856b) from Late Cretaceous Yantardakh amber (Rasnitsyn et al. 2016) was attached with very high probability (97%) to the branch leading to the extant Phygadeuontinae genus *Gelis*. Interestingly, most of the tips to which the Cretaceous fossils attached contained no or only sparse morphological information (Fig. 4a).

**Figure 4.**
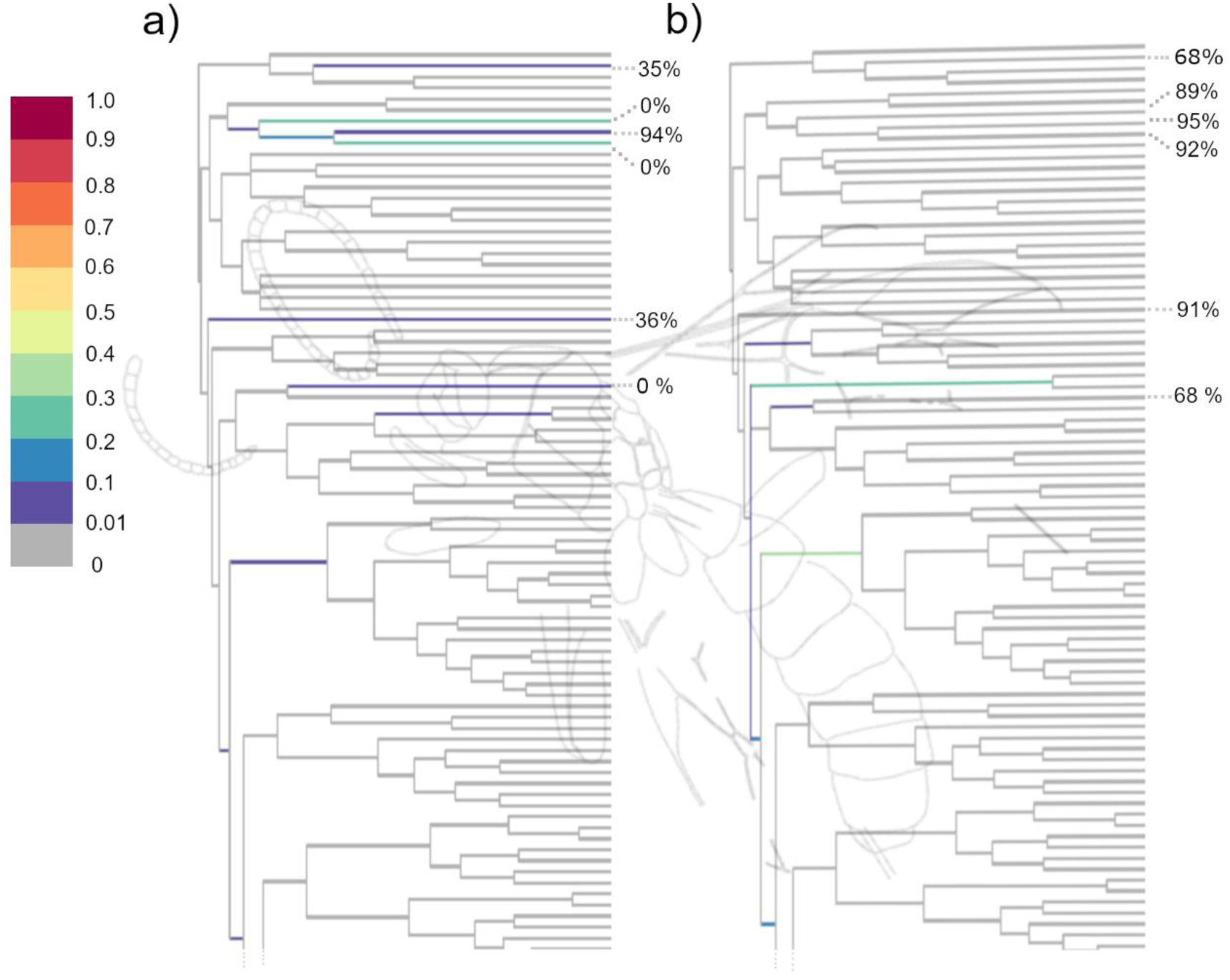
Placement of a representative of the Cretaceous fossils (*Labenopimpla kasparyan*) in the “Xoridinae” analysis a) with and b) without some outgroup taxa missing morphological data. The trees represent majority rule consensus trees with fossils excluded. Branches are colored by probability of a fossil attaching to them (RoguePlots). Percent values refer to the portion of scored morphological characters for a given terminal branch. The remaining Cretaceous compression fossils had similar attachment patterns as the ones depicted here (Supplementary file S9). Background image modified after (Kopylov 2010b).

We repeated the analysis with the “Xoridinae” outgroup setting after improving the scoring of morphological characters for the outgroup taxa, especially those on attracting, bare branches. The placement of the Cretaceous impression fossils shifted considerably and mostly towards the root of the tree compared to the previous analysis (Fig. 4b). Most of the Cretaceous fossils were now placed on stem branches of both non-pimpliform and pimpliform lineages, with some placement probability still on the long branches of outgroup taxa (e.g., *Orthopelma*, *Brachycyrtus*). All Labenopimplinae now attached with the highest probability on the stem branch of the most basal pimpliform subfamily Diplazontinae (23-39%) and on a nearby long branch leading to the extant tryphonine genera *Zagryphus* and *Thymaris* (22-32%). The probabilities of a placement of the Cretaceous impression fossils with crown Banchinae was now close to zero (Fig. 4).

### Impact of fossil placement on age estimates

Although the placement of the Cretaceous fossils significantly changed when the improved morphological matrix was employed, the median age estimates remained relatively stable (Fig. 5). In contrast, there was a more obvious improvement in the precision of the age estimates for all but one of the examined nodes (Supplementary file S8).

**Figure 5.**
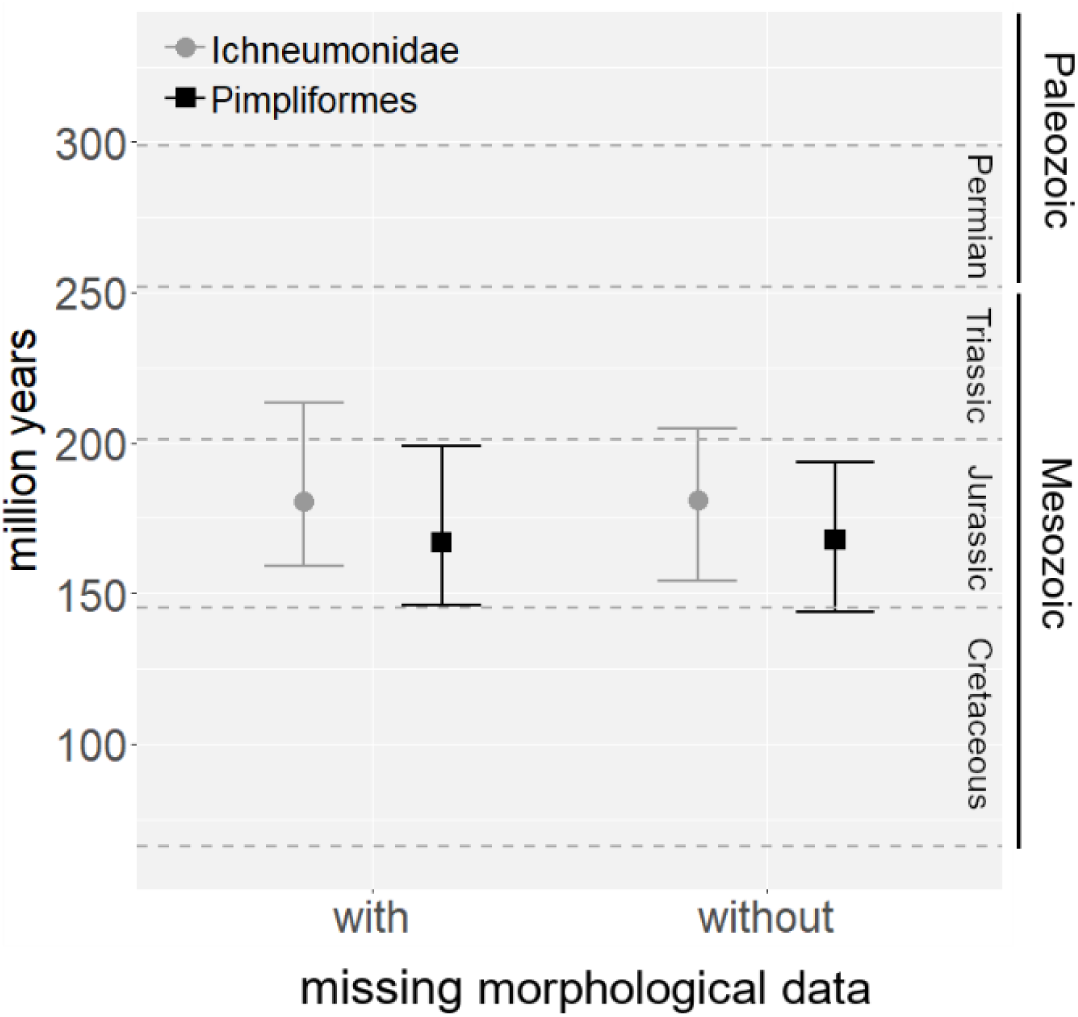
Age estimates in million years for Ichneumonidae and Pimpliformes in “Xoridinae” outgroup analyses with (left) and without (right) “bare-branches” for some outgroup taxa. The median and 95% credibility intervals are plotted on the y axis. Dashed horizontal lines indicate transitions between major geological periods.

### Age of Pimpliformes

The median age estimates for Pimpliformes varied widely across the different outgroup analyses, ranging from 167 Ma to 263 Ma (Fig. 2, Supplementary File S8). We suppose that the age estimates were biased when more distant outgroups were included, as these outgroups were poorly sampled and no fossils provided time information for the outgroup branches to accurately estimate evolutionary rate in this part of the tree. This notion is supported by the smaller clock-rate variance and higher consistency in the age estimates when only close outgroups (“Tanychorinae” and “Xoridinae”) were included.

We thus report here only the preferred age estimates from the “Xoridinae” outgroup setting, with the improved morphological matrix, as we deem initial placement of most of the Cretaceous fossils among extant Banchinae erroneous (Table 2, Fig. 6). The preferred age estimates suggest that Ichneumonidae originated during the Early to Middle Jurassic and Pimpliformes during the Middle to Late Jurassic. The start of the radiation of the extant pimpliform subfamilies is estimated as having occurred in the Early Cretaceous.

**Figure 6.**
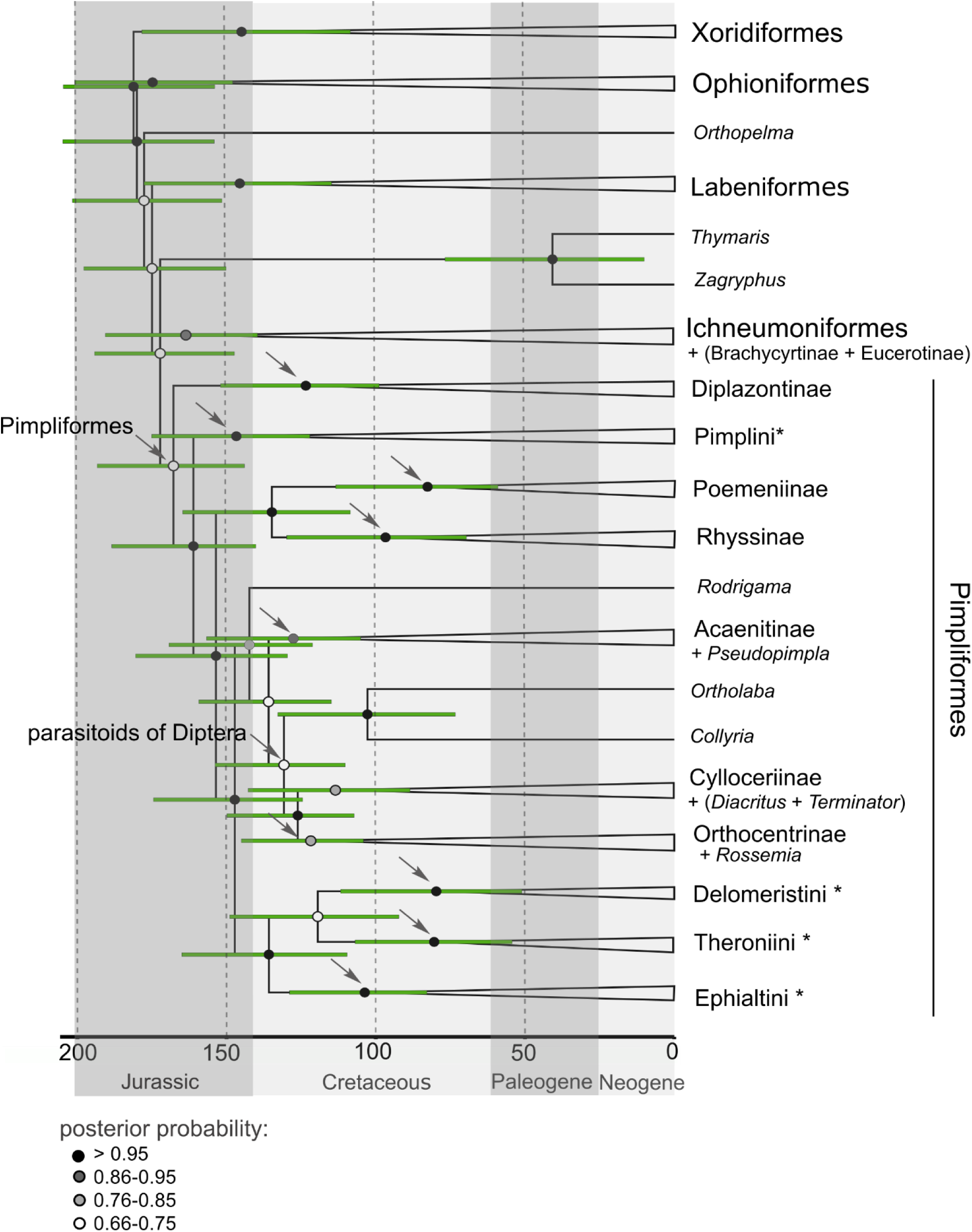
Dated majority rule consensus tree from the total-evidence dating analysis under the “Xoridinae” outgroup setting with improved morphological matrix. The tree contains only extant tips, as the fossils were excluded prior to summarizing tree samples from the MCMC analysis. Most of the clades are collapsed to depict subfamily level relationships among Pimpliformes. Arrows indicate nodes for which age estimates are represented in Table 1. The names of the nodes are given if they do not correspond to the names of their tips. Horizontal bars represent median and 95% credibility intervals for age estimates.

**Table 2.** Age estimates from the preferred analysis with credibility intervals for Ichneumonidae, Pimpliformes and the pimpliform subfamilies. As the subfamily Pimplinae was not recovered as monophyletic, age estimates for the tribes are given (Delomeristini, Ephialtini, Pimplini and Theroniini). The Diptera parasitoids here comprise three subfamilies: Cylloceriinae, Diacritinae and Orthocentrinae (excluding Diplazontinae and the diacritine *Ortholaba*, which did not form a monophyletic group with them).

| **Taxon Group** | **Median** | **Mean** | **95% Credibility** |
| --- | --- | --- | --- |
| Ichneumonidae | 181.2 | 181.4 | 154.0–204.7 |
| Pimpliformes | 167.8 | 168.7 | 144.0–193.4 |
| Diptera parasitoids | 126.2 | 127.8 | 107.3–150.0 |
| Acaenitinae | 127.6 | 129.0 | 105.1–156.7 |
| Diplazontinae | 123.5 | 124.8 | 98.9–152.0 |
| Poemeniinae | 82.7 | 83.5 | 59.1–113.4 |
| Rhyssinae | 96.8 | 97.7 | 69.6–129.9 |
| Delomeristini | 79.7 | 80.6 | 51.1–111.8 |
| Ephialtini | 103.7 | 105.4 | 82.9–128.9 |
| Pimplini | 146.7 | 147.4 | 122.1–175.1 |
| Theroniini | 80.4 | 80.8 | 54.3–106.7 |

## DISCUSSION

### Impact of Outgroup Sampling on Age Estimates

Multiple studies have shown that outgroup choice can greatly affect tree topology estimates, especially in cases with large heterogeneity of branch lengths and uneven taxon sampling (Puslednik and Serb 2008; Ware et al. 2008; Hayes et al. 2009; Thomas et al. 2013; Kirchberger et al. 2014; Wilberg 2015), but its influence on divergence time estimates, especially in the context of total-evidence dating, has scarcely been studied at all. We show here that with the inclusion of more distantly related and/or poorly sampled outgroups, the age estimates become older and often less precise.

In these last six years since total evidence dating became established, a majority of studies have employed rather poor outgroup sampling, typically including one to just a handful of taxa (Ronquist et al. 2012a; Arcila et al. 2015; Dornburg et al. 2015; Close et al. 2016; Herrera and Davalos 2016; Kittel et al. 2016; Lee 2016; Bannikov et al. 2017). We here covered most of the outgroup sampling schemes found in these studies, from including a single taxon from the sister group (“one Braconidae” outgroup setting), over a few taxa from the sister group (“Braconidae”), to the inclusion of a series of more to less closely related outgroup taxa (“full outgroup”).

As in most of the previous studies, our outgroups were not only sparsely sampled for extant taxa, but were also missing fossils and most of the morphological data. All these outgroup settings recovered older and usually less precise age estimates than when restricting our dataset to more closely related outgroup taxa for which greater and more even effort in sampling both of extant and fossil taxa has been applied (“Xoridinae” outgroup setting). Interestingly, when two potentially transitional fossils between Braconidae and Ichneumonidae were used as outgroup (“Tanychorinae” outgroup setting), the median age estimates and clock variance were not affected, but their precision was. This could result from the larger uncertainty in the age of these fossils (Supplementary file S2) and the fact that only very few morphological characters were scored for them (9% and 17%, respectively).

Our results might suggest that many of the previous TED analyses would recover younger and likely more accurate age estimates with more detailed and/or restricted outgroup sampling. Presumably, the effect would be most pronounced for datasets where there is a long, unbroken outgroup branch which introduces large rate heterogeneity among lineages. A similar effect has been demonstrated for node dating in the simulation study by Soares and Schrago (2015), where age estimates were significantly biased when there was a combination of large among lineage rate variation and poor taxon sampling. Both accuracy and precision were affected: the mean age of the node in question was constantly overestimated and precision severely decreased. Large heterogeneity of evolutionary rates can already be identified on a non-clock tree by comparing branch lengths and outgroup taxa accordingly excluded, as we exemplified in our study; but this has rarely been employed in total evidence dating studies (but see Grimm et al. 2015).

### Challenges for fossil placement in TED

In TED analyses, the placement of fossils is solely dependent on the available morphological information for both fossil and extant taxa. The quality of fossil placement based on morphological matrices is thus primarily limited by imperfect preservation of fossils, but also by high levels of morphological homoplasy, which has been reported for ichneumonids (Gauld and Mound 1982; Klopfstein and Spasojevic 2019). We here identified another potentially major issue in TED studies, which we called “bare-branch attraction”: the tendency of especially older fossils to attach to terminal branches leading to extant taxa for which no morphological data has been collected. This artefact exposes the dangers of insufficient sampling of extant taxa for morphology, which can distort fossil placement and as a consequence age estimates in TED analyses.

Many of the previous TED studies included from a few to more than half of the extant taxa without morphological data, while some of the extant taxa scored for morphology had high amounts of missing data (Ronquist et al. 2012b; Arcila et al. 2015; Dornburg et al. 2015; Harrington and Reeder 2017). Guillerme and Cooper (2016) addressed this issue in the context of topology reconstruction in TED analyses. In their simulations, topology estimates were more negatively affected by a large percentage of extant taxa with missing morphological data than by any other analyzed parameter. It remains unclear to what extent their results were influenced by bare-branch attraction, as they did not analyze individual fossil placements, but the artefact was likely playing up under their scenario as well.

In our study, bare-branch attraction mostly concerned the compression fossils from the Cretaceous. These fossils are all, expect *T. xoridoptera*, classified in two extinct subfamilies, Palaeoichneumoninae and Labenopimplinae. The phylogenetic position of these subfamilies is unclear, but two options have been suggested: a transitional position between Tanychorinae and extant Ichneumonidae, which would mean they represent stem ichneumonids, or some rather basal position as crown ichneumonids (Kopylov 2009, 2010b). In fact, the name “Labenopimplinae” reflects some similarity to the extant subfamilies Labeninae and Pimplinae, which would indicate the latter option (Kopylov 2010b).

We showed that the predominant placement of most Labenopimplinae (and other Cretaceous compression fossils) with crown group Banchinae and Tersilochinae in our initial analysis, was a consequence of the bare-branch attraction artefact. Among the branches where these fossils attached on the tree, only a single tip (*Apophua*) contained morphological data, while the remaining taxa were only sampled for molecular characters. Some other, younger fossils with rather diffuse placement also often attached to these branches, which might suggest that when morphological information for the placement of a fossil is limited, TED analyses tend to place a fossil on “bare-branches”, especially if those branches are long. Decreasing the percentage of extant taxa with missing morphological information, especially among the outgroups, from 27% to 16% was enough to reverse the initially erroneous placements of the oldest included compression fossils, which were instead placed on rather basal branches of crown group ichneumonids, in accordance with the second hypothesis suggested at the time of their original description (Kopylov 2009, 2010b).

It remains to be shown whether older and/or ancestral fossils are more prone to bare-branch attraction, or if it influences fossils of all ages to a similar extent. Additional analyses are also needed to determine under which conditions the erroneous fossil placement is leading to biased age estimates. In our study, we did not see a large negative impact of bare-branch attraction, but this will presumably not always be the case – we might just have been lucky that the attracting bare branches were spanning the age of the true placements as well, which was then informed correctly by the numerous remaining fossils. And even in our case, the precision of age estimates was improved by reducing the number of bare branches in the phylogeny.

### The age of Pimpliformes and the biological context of their diversification

Our most credible analysis estimated the median age of the family Ichneumonidae to 184 Ma (95% CI: 168.3–203.9 Ma) and of Pimpliformes to 172 Ma (95% CI: 157.3–191.1 Ma), i.e., the Early and Middle Jurassic period, respectively. Compared to the fossil record, the estimated age for Ichneumonidae is 60–70 Ma older than the oldest certain ichneumonid fossils, which would imply a rather long ghost lineage. However, a similar gap in the fossil record is present between the oldest and the second oldest ichneumonid fossils (Palaeoichneumoninae and Labenopimplinae: between 26 and 53 Ma, depending on the adopted age of the geological formations in question), suggesting that the implied ghost range is not that long after all. Such a gap is even more acceptable if we consider the paucity of Jurassic Hymenoptera fossils in general (Rasnitsyn and Quicke 2002). Furthermore, the last few years have seen unexpected discoveries of fossils that have closed large gaps between much older (molecular) age estimates and the previously known fossil record, for instance in Lepidoptera (Eldijk et al. 2018).

Further insights into the age of ichneumonids come from age estimates for their hosts, predominantly holometabolous insects, which had to originate before their parasitoids could radiate. Initially, the radiations of the biggest orders of holometabolous insects, Hymenoptera, Coleoptera, Diptera and Lepidoptera, were believed to have been associated with the radiation of flowering plants and were dated to the Early Cretaceous (Grimaldi 1999; Misof et al. 2014). However, these estimates were later deemed too young, and a Late Permian origin was suggested based on a re-analysis of a large phylogenomic dataset (Misof et al. 2014) with more appropriate calibration points (Tong et al. 2015). This later study implies that the major host groups for ichneumonids were already present during the Jurassic, when both Ichneumonidae and Pimpliformes originated, suggesting that the radiation of these parasitoids might have happened only shortly after the radiation of their host groups.

### Taxonomic implications

To date, some relationships among pimpliform subfamilies have remained unresolved, most likely due to a rapid radiation early in their evolution (Klopfstein et al. 2019). In our study, we utilized a small set of eleven genes but with an extensive sampling of pimpliform genera and morphological characters. We recovered a very similar tree topology as in the nucleotide versions of the previous analysis. The few differences mainly concern relationships within the clade of koinobiont endoparasitoids, but these were weakly resolved in both analyses. However, with our increased taxon sampling, we recover some previously unclear or unidentified relationships.

The monophyly of Acaenitinae was never supported in previous molecular analyses (Quicke et al. 2009; Klopfstein et al. 2019), while we recovered it with some support (posterior probability around 0.8), but including the Ephialtini genus *Pseudopimpla*. Acaenitines are morphologically easily recognized by the large and elongate hypopygium, but are otherwise a rather heterogenous group (Förster 1869; Broad et al. 2018). Although the hypopygium of *Pseudopimpla* is not as large as in *Coleocentrus*, with which it clustered in our analyses, it is strongly sclerotized and clearly elongate, as in some other acaenitine genera (e.g., *Procinetus, Leptacaenitus*, *Prosacron*). In addition, *Pseudopimpla* also has a very elongate eighth tergite in females, as is often seen in Acaenitinae (including *Coleocentrus*) and only rarely in Pimplinae (*Pimplaetus*, *Pachymelos*); the eighth tergite is also elongate in males of *Pseudopimpla*, as well as in *Coleocentrus*. The presence of lobes on the tarsal claws in *Pseudopimpla* suggests a placement of the genus in Ephialtini (Gauld et al. 2002), but these lobes have evolved several times independently in ichneumonids in any case (at least in some Pimplinae, Labeninae, Orthopelmatinae and Collyriinae), and are also present in the acaenitine genus *Hallocinetus* (Townes 1971). It is noteworthy that *Pseudopimpla* lacks one of the apomorphies used to define Pimplinae, namely a sculpturally differentiated posterior band on the metasomal tergites. We thus here transfer *Pseudopimpla* to Acaenitinae.

The placements of the cyllocerine *Allomacrus* and orthocentrine *Terminator* with *Diacritus*, and of the cylloceriine *Rossemia* within Orthocentrinae are also strongly supported, but we refrain from any taxonomic changes, considering the poor resolution at subfamily level in this part of the tree. In any case, our results might suggest that Cylloceriinae, Diacritinae and Orthocentrinae constitute an entity which is difficult to subdivide, based on both molecular and morphological evidence. The close association of these three subfamilies is also supported by their biology, where known, as koinobiont endoparasitoids of Diptera, so these three subfamilies might be synonymized in the future. However, there are no reliable host records for *Allomacrus*, *Diacritus*, *Ortholaba*, *Rossemia* or *Terminator* (Broad et al. 2018), and we thus refrain here from any formal changes.

The monophyly of the subfamily Pimplinae has already been questioned in the past. It has been supported both with the morphological (Gauld et al. 2002) and the latest 28S dataset (Quicke et al. 2009), but some earlier analyses had recovered Pimplini separately from Ephialtini (Belshaw et al. 1998; Quicke et al. 2000). This was also the case in Klopfstein et al. (2019), where at least the genus *Xanthopimpla* (in nucleotide and amino acid analyses), if not the entire Pimplini (in nucleotide analyses), clustered apart from the remaining Pimplinae. Morphologically, there are not many clear synapomorphies for Pimplinae (Gauld et al. 2002), and the Pimplini show many plesiomorphic character states, such as a quadrate areolet, rather stout and short first tergite with dorsal median carinae, and an ovipositor of intermediate length, all of which have had already been present in some Cretaceous Labenopimplinae and could also have had been present at the base of Pimpliformes. Further investigation, such as the detection of possible anomalies in the nucleotide data, is needed in order to decide whether *Xanthopimpla* and *Lissopimpla* or even all of Pimplini should be transferred to a separate subfamily.

## CONCLUSIONS

We have demonstrated that poor outgroup sampling can negatively affect both accuracy and precision of age estimates in total-evidence dating analyses. To achieve more reliable age estimates, one should consider a more detailed taxon sampling that includes morphological and fossil data from a closely related outgroup, or alternatively first perform a non-clock analysis including outgroups, and then use topology constraints to correctly position the root in the dating analysis, while excluding any sparsely sampled outgroup taxa.

We also illustrated the importance of careful consideration of fossil placement in total-evidence dating analyses in order to identify artefacts or biases. The bare-branch attraction artefact that we have discovered here might turn out to be universally problematic for TED. Thus, it deserves further assessment in the future, possibly through simulation studies, but it can easily be circumvented by a more complete sampling of morphology for extant taxa.

Finally, we provided the first age estimate for the extremely diverse group of ichneumonid parasitoid wasps. The obtained Jurassic origin for the family and for Pimpliformes agrees with the timing of the radiation of their major host groups. It remains to be seen how the age estimate for the family will change when more fossils are included in a TED analysis; this will require a major upgrade of the morphological matrix that includes filling in all the missing morphological information and improving the sampling of non-pimpliform ichneumonid extant taxa.

## AKNOWLEDGEMENTS

For access to fossil specimens, we are grateful to Marsh Finnegan and Alan M. Rulis (Smithsonian National Museum of Natural History, Washington, USA), Talia Karim and David Zelagin (Museum of Natural History, University of Colorado, Boulder, USA) and Ricardo Pérez-de la Fuente (Museum of Comparative Zoology, Harvard University, USA). We also thank Dmitry Kopylov (Borissiak Paleontological Institute, Russian Academy of Sciences, Moscow, Russia) and Sonja Wedmann (Forschungsstation Grube Messel, Senckenberg, Germany) for providing access to additional fossil specimens and for commenting on derived fossil ages. We are grateful to Eric Chapman and Mike Sharkey (Kentucky, USA), Rikio Matsumoto (Osaka, Japan), and the Swedish Malaise Trap project for extensive material for morphological and molecular studies. Remo Ryser (German Centre for Integrative Biodiversity Research, Leipzig, Germany) and Djordje Markovic (Institute of Zoology, University of Belgrade, Belgrade, Serbia) contributed to the lab work.

## SUPPLEMENTARY MATERIAL

Data available from the Dryad Digital Repository: http://dx.doi.org/10.5061/dryad.[NNNN]

